# Genetic mapping of the powdery mildew resistance gene Pm13 on oat chromosome 1D

**DOI:** 10.1101/2024.01.26.576824

**Authors:** Selma Schurack, Steffen Beuch, Sandy Cowan, Irene Griffiths, Magdalena Lunzer, Laura Morales, Sara Tudor, Hermann Buerstmayr, Catherine J Howarth, Nicholas A Tinker, Matthias H Herrmann

**Affiliations:** Julius Kühn Institute (JKI), Federal Research Centre for Cultivated Plants, Institute for Breeding Research on Agricultural Crops, Rudolf-Schick-Platz 3a, 18190 Sanitz, Germany; Nordsaat Saatzucht GmbH, Saatzucht Granskevitz, Granskevitz 3, 18569 Schaprode, Germany; Institute of Biological, Environmental and Rural Sciences, Aberystwyth University, Aberystwyth, SY23 3EE, UK; Department of Agrobiotechnology Tulln, BOKU-University of Natural Resources and Life Sciences Vienna, Konrad Lorenz Str. 20, 3430 Tulln, Austria; Agriculture and Agri-Food Canada, Ottawa Research and Development Centre, 960 Carling Ave., Ottawa, ON K1A 0C6, Canada

**Keywords:** Oats, *Avena sativa*, *Blumeria graminis*, powdery mildew resistance, adult plant resistance, GWAS, QTL

## Abstract

Powdery mildew, caused by the biotrophic fungus *Blumeria graminis* DC. f. sp. *avenae*, is a widespread disease of oats, especially in the temperate regions of Western and Central Europe, and the use of resistant varieties is the most sustainable way to ensure stable yields. Therefore, the identification of robust and effective resistance to powdery mildew is of great interest for oat breeding. In contrast to race-specific resistance genes, adult plant resistance (APR) is generally considered to be more durable. The oat variety Firth, as well as related varieties such as Husky or Flämingstip, contains an unknown APR gene, which was previously located on chromosome 1D using DArT markers. The aim of this study was to confirm and refine the chromosomal location of this resistance gene, tentatively named Pm13. To this end, two independent experiments were carried out using different genetic material under natural infection conditions in the field: genome-wide association mapping (GWAS) in a diverse set of 250 oat lines grown in ten environments and QTL mapping in a HuskyxAVE1284 bi-parental population grown in three environments. Both approaches identified a QTL for powdery mildew resistance on the distal end of chromosome 1D in the hexaploid Sang oat genome. The locus explained up to 15 % of the phenotypic variance in GWAS and 64 % of the phenotypic variance in QTL mapping. Comparison of field data with results from laboratory leaf segment tests confirmed that Pm13 does indeed confer APR. The sequence information of the identified linked markers may allow the development of molecular markers useful for early selection of oat lines with high levels of APR.

## Introduction

Cultivated oats (*Avena sativa* L.) are an important food and feed crop in many countries. The largest production areas are in Russia, Canada, Australia, Brazil, China and the USA. Despite the increasing demand for milling oats, the production volume of oats in Europe has fluctuated between 9730 MT (2002) and 6940 MT (2018)^1^ since 1999. This production volume represents around 2-3% of total cereal production in Europe. Oats can therefore be described as a minor crop, with a somewhat higher importance in northern countries such as Sweden, Finland, Norway and the UK. With increasing prosperity and a greater focus on quality and nutrition, oats are receiving more attention in research, production, processing and product development. However, production is affected by several pathogens, including *Puccina coronata* f. sp. *avenae* and *Blumeria graminis* f. sp. *avenae*, and abiotic stresses such as drought.

Oat powdery mildew (PM) is caused by the obligate biotrophic fungus *Blumeria graminis* f. sp. *avenae* (BGA). This oat pathogen is mainly found in temperate areas of Europe, but also in Mediterranean countries such as Spain, Italy and Morocco. The economic importance of BGA correlates with the management status of the crop. Dense stands with a good nitrogen supply allow stronger PM development. In addition, yield losses due to BGA depend on weather conditions and the time of infection and can be significant in susceptible cultivars (Roderick et al., 2000). Resistance breeding against PM has been quite successful in Europe over the last 30 years. Many used and unused sources of resistance within the different *Avena* species have been described by Jones and Griffiths (1951), Hayes and Jones (1966), Hoppe and Kummer (1991), Herrmann and Roderick (1996), Hsam et al. (1997, 1998), Sánchez-Martín et al. (2011), Okoń et al. (2014, 2016, 2018, 2021), Okoń and Ociepa (2018) and Okoń and Kowalczyk (2020). So far, 13 BGA resistance genes have been catalogued and genetically mapped in oats (Admassu-Yimer et al., 2022; Herrmann and Mohler, 2018; Hsam et al., 2014; Ociepa et al., 2020; Ociepa and Okón, 2022). Most are effective at both the adult and seedling stages, but some resistances are only effective at the adult stage(Hite et al., 1977; Roderick et al., 1995, 1991). Two sources of resistance widely used in breeding have been effective in Europe for decades. The first is the resistance in the crown rust differential line Pc54, described by Sebesta et al. (1993) and recently mapped to chromosome 1A by Admassu-Yimer et al. (2022). The second adult plant resistance was developed by Lochow-Petkus GmbH (Germany) with the cultivars Firth and Winston, two sister lines with good APR to BGA and incomplete resistance to crown rust and smut. These varieties have been used as parents in several crossing programmes, resulting in varieties such as Flämingstip and Husky. A phylogenetic tree based on 8417 GBS markers (Brodführer et al., 2023) reflects the relationship between Husky, Flämingstip, Firth, Winston and some breeding lines (Supplementary Figure S1). Regarding the APR in Firth, a first QTL mapping approach was reported by Hagmann et al. (2012), who described a major QTL near the DArT marker oPt-6125, which was mapped to linkage group 5_30 on the Kanota/Ogle map (KxO, Tinker et al., 2009). Hagmann et al. (2012) found a continuous distribution of infection levels under field conditions across the whole range for the 184 RILs studied. The KxO linkage group 5_30 was later assigned to chromosome 1D (Chaffin et al., 2016). Based on known pedigree and GBS data, we hypothesised that the APR in Firth, Flämingstip and Husky has the same genetic background. To verify this assumption and to obtain a more precise localisation of the corresponding QTL, we performed two independent experiments, one with Husky (QTL mapping with bi-parental population) and one with Firth and related breeding lines (GWAS with FO-panel).

## Materials and Methods

### Plant Materials

For GWAS, a facultative oat panel (FO-panel) with 250 lines of similar panicle emergence and plant height of the larger KLAR panel described in Brodführer et al. (2023) was used. For QTL mapping, a HuskyxAVE1284 bi-parental population was developed. The initial cross between the resistant variety Husky and the susceptible landrace AVE1284 was followed by a backcross of F_1_-plants to Husky and selfing of two BC_1_ individuals for development of RILs. The resulting population comprised 176 individuals, divided into two subpopulations of 101 (subpopulation 1) and 75 (subpopulation 2) individuals.

### Field Trials and PM scoring

For GWAS, the FO-panel was grown in an augmented design including ten checks with five replications in field trials located in Aberystwyth, United Kingdom (UAB), Granskevitz, Germany (NORD) and Tulln, Austria (BOKU). At each location, the experiments were sown in the autumn (AU) and spring (SP) in 4 to 6 m^2^ plots. For UAB and BOKU, the experiment was performed in 2022 and 2023; for NORD the experiment was performed in 2022, resulting in ten location:sowing-season:year environments (UAB_AU22, UAB_SP22, UAB_AU23, UAB_SP23, BOKU_AU22, BOKU_SP22, BOKU_SP23, BOKU_AU23, NORD_SP22, NORD_AU22). Best Linear Unbiased Estimators (BLUEs) were estimated with the mmer function from the sommer R package (Covarrubias-Pazaran, 2016) with row and column effects set as random. Because of the augmented design without replications and the uncertainties of scoring a plot without repetitions, GWAS was not performed for environments individually, but across all environments.

For QTL mapping, the HuskyxAVE1284 population was grown in a randomised complete block design with two replications in Groß Lüsewitz, Germany (GL) in 2021 and 2022 and Quedlinburg, Germany (QLB) in 2021. A plot consisted of a double row of 1 m length and 20 cm distance between rows and plots.

In both approaches, the severity of natural powdery mildew infection of the leaves was scored as percent infection [%] or on a 1-9 scale. Scorings were converted to percent for data analysis using the following formula: p_z_ = (maxp_z_ -minp_z_)/2+minp_z_; with p_z_ being the converted percent value of score z, minp_z_ being the lower percent value of the percent value range attributed to score z and maxp_z_ being the maximum percent value of the range attributed to score z.

### Detached leaf segment test

In addition to nursery experiments, a detached leaf segment test comprising 360 entries from the entire KLAR panel was performed. Plants were grown in pots in a glasshouse under controlled temperature conditions (20 ± 3 °C during the day and 16 ± 3 °C during the night). Following the procedure described in Herrmann and Mohler (2018), leaf segments from three 14-day-old seedlings per entry were deposited on benzimidazole agar. The leaf segments were inoculated with freshly harvested spores using an infection tower. A powdery mildew isolate found in Groß Lüsewitz, Germany with virulence for Pm1, Pm3, Pm6 but no virulence for Pm9, Pm5, and Pm7, based on the reaction of differentials involved in the experiment, was used. Eight to ten days after inoculation, the percentage of leaf covered with powdery mildew was estimated for each leaf segment.

### Statistical Analysis

In the ANOVA with BLUEs of the FO-panel restricted to the replicated standards, a complete model with all factors (all random: environments = 5; seasons = 2, genotypes = 10, replications = 6) and all interactions was used. For the ANOVA with BLUEs of the unreplicated entries, a complete model with environments (5), seasons (2), and genotypes (250) defined as random factors, and with the respective interactions was used in PLABSTAT (Utz, 2011). PM severity data of the HuskyxAVE1284 tests at three environments were transformed via the logit function and analysed via an ANOVA and a heritability estimation in PLABSTAT (Utz, 2011) defining environments, replication and genotype as random in the model.

### GWAS

Molecular marker data of the 250 oat lines in the FO panel was extracted from Brodführer et al., (2023). The Genomic Association and Prediction Integrated Tool (GAPIT3) package in R (Version 4.1.0) was used for GWAS analysis (Wang et al., 2021). GWAS was performed including three PCs with four different models that all yielded reasonable QQ plots (Blink, FarmCPU, MLM, MLMM) and with a minor allele frequency cut-off of > 0.05. A Bonferroni multiple test threshold was used to determine significance. For analysis of candidate genes, the significant region was extended to the next non-significant marker downstream and upstream of a significant marker-trait association. For investigation of the relationship of the oat lines, molecular marker data extracted from Brodführer et al. (2023) was used in TASSEL to construct a phylogenetic tree. Markers were filtered for a minor allele frequency of 0.05 and 95 % completeness.

### QTL mapping

Genomic DNA of 176 individuals of the HuskyxAVE1284 population was isolated from fresh leaves of a bulk of five plants according to the CTAB procedure described by (Stein et al., 2001). A GBS procedure followed by a capture-based complexity reduction (Rapture) was applied to increase the depth of sequencing for a restricted and consistent set of GBS fragments. Rapture-based GBS-libraries were prepared by the Genomic Analysis Platform, Institute of Integrative Biology and Systems, Laval University (Québec, Canada) according to Bekele et al. (2020). Pooled, barcoded libraries were sequenced at the Génome Québec Innovation Centre (Montreal, Canada) on a single lane of a NovaSeq 6000 sequencer (Illumina, San Diego, CA). GBS analysis was performed as described by Bekele et al. (2018). Briefly: raw reads in fastq format were deconvoluted into a single tag-count file per entry using the software UNEAK (Lu et al., 2013). Tag count files were then analysed using the production mode of the software Haplotag (Tinker et al., 2016) based on a tag nomenclature established by Bekele et al., (2018). This procedure results in variant calls for tag-level haplotypes having a standardised nomenclature that was later positioned on the Sang reference genome (Kamal et al., 2022) as described by Tinker et al. (2022). Variant calls for tag-level haplotypes were filtered to keep those with a minimum 50% data completeness across progeny. These variants were ordered based on the Sang genome and phased based on parental alleles. Missing data were imputed using a script described by Tinker et al., (2022). This algorithm drops markers that are unlikely to be ordered correctly based on double crossovers, and imputes remaining markers based on genotypes of flanking markers. Genetic maps were constructed for both subpopulations individually in JoinMap®4.1 (van Ooijen, 2006) with 1488 loci. Identical loci and those with more than 30 % missing data were excluded (687 and 470 loci remaining for subpopulation 1 and 2, respectively). Linkage groups were created with pairwise recombination frequency and then maps were calculated with the Kosambi function using a LOD threshold of 1.00 and a recombination threshold of 0.4.

QTL detection was performed using Haley-Knott regression with the R package R/qtl v1.46-2 (Broman et al., 2003). The cross type was set to “bcsft”. LOD thresholds to detect QTL were determined by 1000 permutations in the scanone function and the global type I error was set to 5 %. The QTL LOD support interval was extended to the nearest neighbouring marker or chromosome end. The percentage of explained variance (PVE) was estimated with the following formula: PVE = 1 – 10^((-2*LOD) / n)^, where n equals the sample size, and LOD is the LOD peak value.

Linkage disequilibrium was analysed in TASSEL. The naming of the QTLs follows the latest nomenclature for oat genes (Jellen et al., 2024).

## Results

### GWAS

To investigate genetic loci associated with PM resistance, PM severity was scored in autumn- and spring-sown trials of the FO-panel. Figure 1 shows the observed PM severities for all environments. Strikingly, PM severity was significantly lower in the autumn-sown trials than in the spring-sown trials. PM resistance showed reproducible differences among the tested oat panel across years, locations and sowing seasons, as indicated by an estimated broad-sense heritability of 89 % for the replicated standards and 83 % for the unreplicated entries (Supplementary table S1).

**Figure 1.**
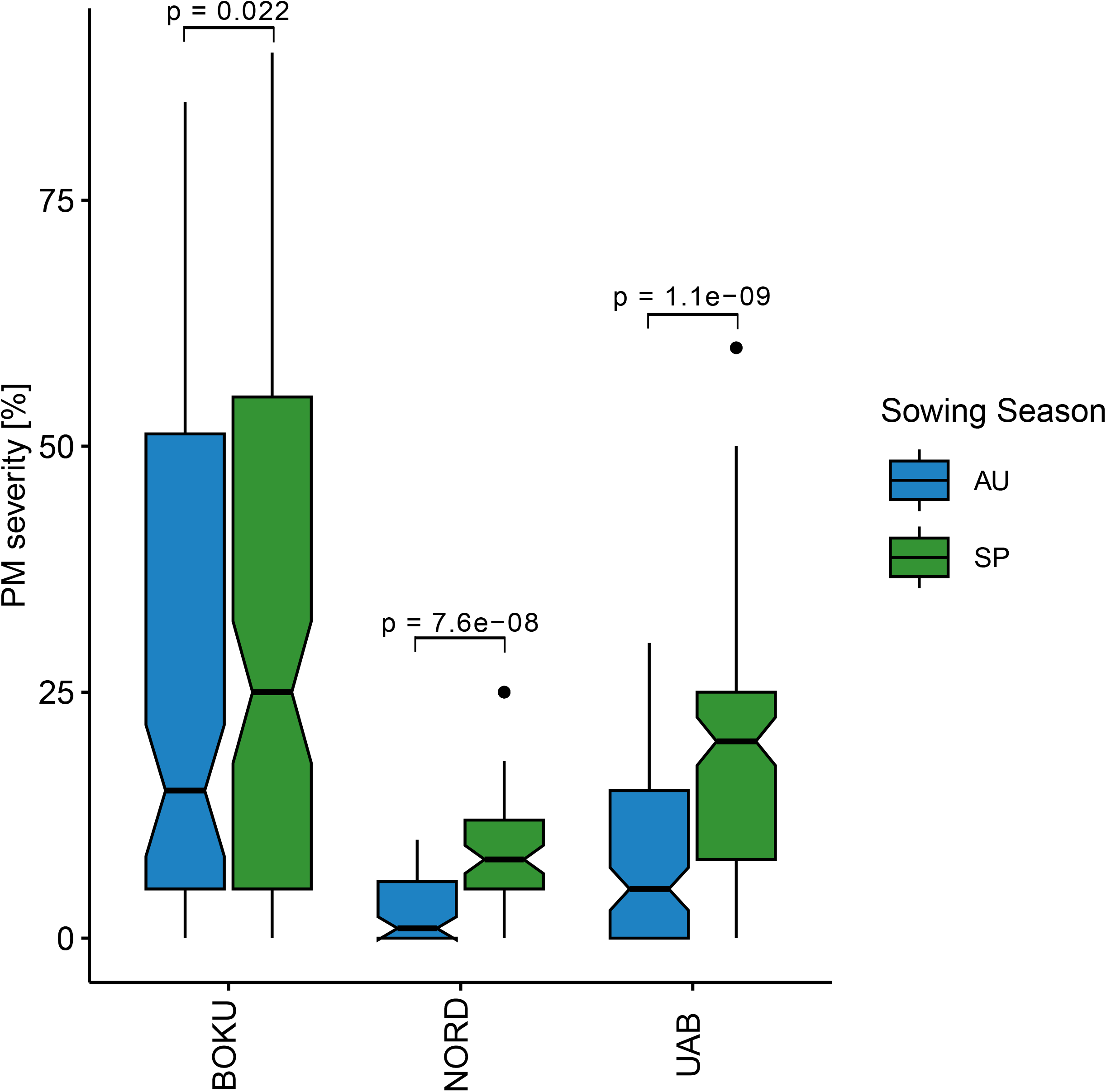
Powdery mildew severity [%] in the FO-panel across years for the different locations and sowing seasons. Kruskal-Wallis p values are given above the brackets. AU: Autumn, SP: Spring.

Principal component analysis (PCA) of the genotypes in the FO-panel showed a relatively homogenous population structure (Figure 2B). GWAS was performed for the BLUEs for PM severity across all environments with different statistical models that all showed acceptable QQ plots (MLM, MLMM, FarmCPU, BLINK; Figure 2C). Overall, thirteen statistically significant marker-trait associations (MTAs) located on six chromosomes were identified by the different models (Figure 2D, Table 1). Four of the MTAs were located on the distal end of chromosome 5D, where the highly effective PM resistance gene Pm7 is positioned (Brodführer et al., 2023). Another four MTAs were located on 1D, the chromosome the APR gene from Firth was assigned to previously (Hagmann et al., 2012). The phenotypic variance explained by the MTAs identified on 1D ranged from 1.12 % (avgbs_cluster_38613.1.56) to 15.25 % (avgbs2_84146.1.21, Table 1, Supplementary Table S2), the phenotype distributions are presented in Figure 2E. One of the MTAs on 1D was located at position 343715080 and the remaining three MTAs were located at the distal end between positions 418134649 and 440373230, resulting in three significant regions (QPM.FO_1D.1, QPM.FO_1D.2, QPM.FO_1D.3). Significant regions on 1D spanned 0.46 – 1.04 Mb and contained ten to 24 high-confidence genes based on the Sang genome assembly (Kamal et al., 2022). Interestingly, two of the identified markers (avgbs2_84146.1.21 and avgbs_47533.1.46) were located at different positions of the same gene AVESA.00010b.r2.1DG0136610.1, which is predicted to encode a protein kinase superfamily protein. Protein kinases play a crucial role in signal transduction processes and have been proposed to play important roles in quantitative/adult plant resistance (Delplace et al., 2020). Furthermore, avgbs2_84146.1.21 explained the highest proportion of phenotypic variance within the markers identified on chromosome 1D (Table 1), making this locus the most promising candidate. In addition to MTAs on 1D and 5D, BLINK and FarmCPU GWAS models identified MTAs on 1A, 6C, 7A and 7D (Table 1).

**Table 1.** SNP markers associated with powdery mildew severity from GWAS using different models. Chr.: Chromosome. Pos.: Position in the Sang genome assembly. MAF: Minor allele frequency. PVE: Phenotypic variance explained [%].

| SNP | QTL | Chr | Pos | MAF | P.value<br>BLINK | P.value<br>FarmCPU | P.value<br>MLMM | P.value<br>MLM | PVE<br>BLINK | PVE<br>FarmCPU | PVE<br>MLMM | PVE<br>MLM |
| --- | --- | --- | --- | --- | --- | --- | --- | --- | --- | --- | --- | --- |
| avgbs2_89610.1.54 | QPM.FO_1A | 1A | 380816519 | 0.18 | ns | 5.20E-07 | ns | ns |  | 3.75 |  |  |
| avgbs_cluster_38613.1.56 | QPM.FO_1D.1 | 1D | 343715080 | 0.48 | 4.78E-08 | ns | ns | ns | 1.12 |  |  |  |
| avgbs2_84146.1.21 | QPM.FO_1D.2 | 1D | 418134649 | 0.07 | 4.90E-08 | ns | 3.20E-06 | 3.20E-06 | 5.80 |  | 15.25 | 15.25 |
| avgbs_47533.1.46 | QPM.FO_1D.2 | 1D | 418137530 | 0.45 | ns | 9.34E-07 | ns | ns |  | 1.89 |  |  |
| avgbs_69900.1.31 | QPM.FO_1D.3 | 1D | 440373230 | 0.31 | ns | 3.23E-09 | ns | ns |  | 3.51 |  |  |
| avgbs_80265.1.22 | QPM.FO_5D.1 | 5D | 464707770 | 0.16 | ns | ns | 2.75E-09 | 2.75E-09 |  |  | 9.16 | 9.16 |
| avgbs_40526.1.18 | QPM.FO_5D.2 | 5D | 465063194 | 0.15 | 1.94E-21 | ns | ns | ns | 6.65 |  |  |  |
| avgbs_cluster_4805.1.42 | QPM.FO_5D.3 | 5D | 476948334 | 0.06 | 8.55E-07 | ns | ns | ns | 6.81 |  |  |  |
| avgbs_102450.1.35 | QPM.FO_5D.4 | 5D | 483446385 | 0.10 | 2.38E-38 | 2.26E-27 | 2.04E-25 | 2.04E-25 | 18.00 | 28.83 | 34.97 | 34.97 |
| avgbs_cluster_12340.1.33 | QPM.FO_6C.1 | 6C | 535970540 | 0.28 | 7.55E-11 | ns | ns | ns | 0.97 |  |  |  |
| avgbs_97815.1.55 | QPM.FO_6C.2 | 6C | 100827967 | 0.20 | 1.27E-06 | ns | ns | ns | 2.10 |  |  |  |
| avgbs_237792.1.17 | QPM.FO_7A | 7A | 365214647 | 0.11 | 3.58E-11 | ns | ns | ns | 2.20 |  |  |  |

**Figure 2.**
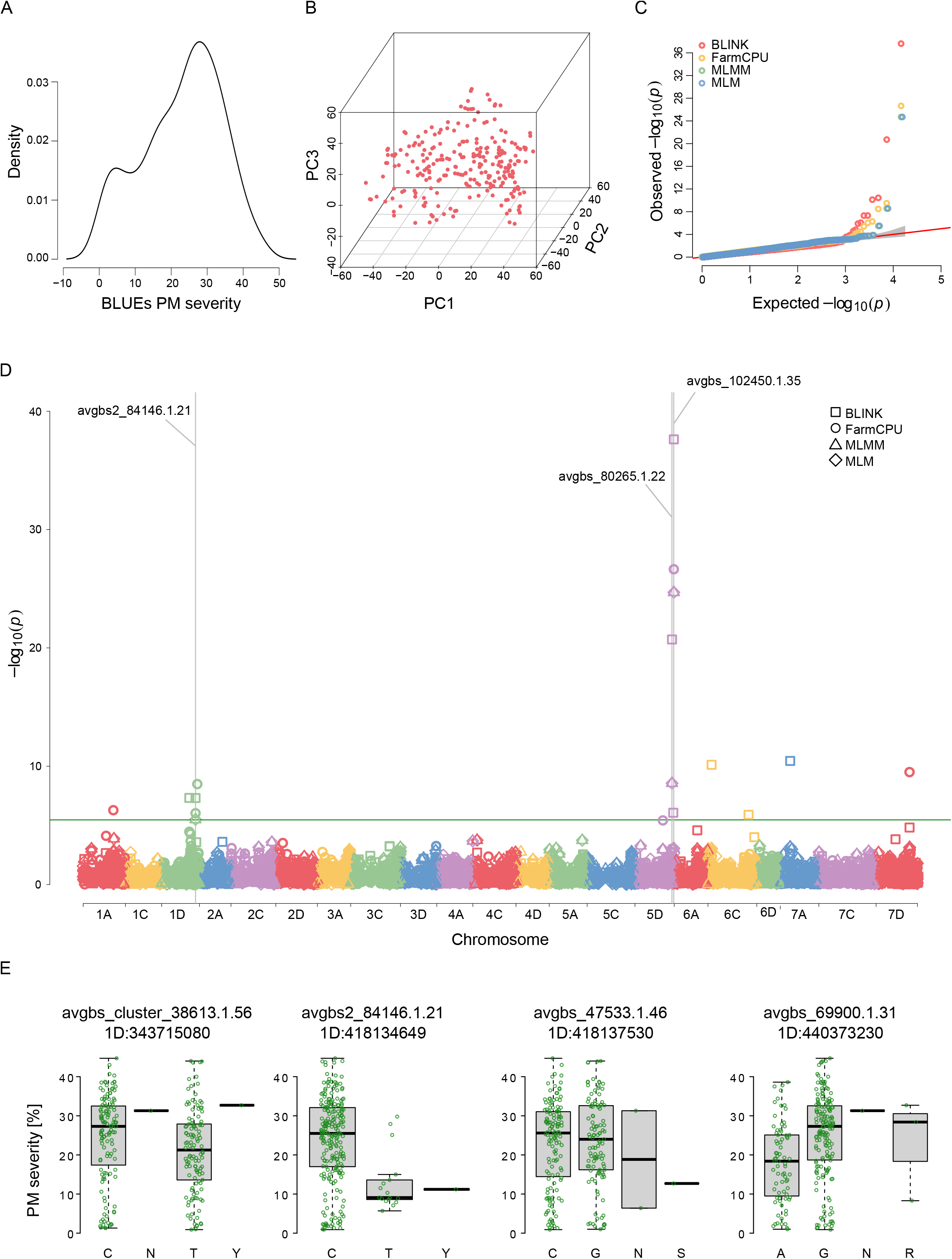
GWAS for powdery mildew severity. **A**. Distribution of the Best Linear Unbiased Estimations (BLUEs) for powdery mildew severity [%]. **B**. Principal component analysis of the investigated genotypes. PC1, PC2 and PC3 refer to the first, second and third components, respectively. **C**. Quantile-quantile (QQ) plot of expected vs. observed p-values of the different GWAS models. The grey area shows the 95% confidence interval. **D**. Manhattan plot of -Log_10_(p) vs. the chromosomal position of SNP markers associated with powdery mildew severity. The solid green horizontal line represents the Bonferroni-corrected p-value cut-off. The grey vertical lines highlight common significant SNP markers detected by at least two models. **E**. Phenotype distribution of the genotypes for significant SNP markers detected on chromosome 1D. Marker names with their respective chromosomal position based on the Sang genome assembly are given at the top. Genotypes are given in IUPAC code (Johnson, 2010).

To verify whether the identified locus indeed confers APR, a detached leaf segment test (LST) was conducted. For this, seedlings of oat lines from the GWAS population carrying the putative resistance allele at avgbs2_84146.1.21 (QPM.FO_1D.2), the marker that explained the largest proportion of phenotypic variance for 1D and that was identified by multiple models, were used. For contrast, oat lines carrying the plant stage-independent resistance gene Pm7 (QPM.FO_5D.4; favourable allele at marker avgbs_102450.1.35) were included as well and results were compared to the field infections of the GWAS experiments. For QPM.FO_1D.2, mean PM severity was significantly higher in the LST than in the field, whereas for QPM.FO_5D.4, PM severity was very low in LST as well as in the field (Figure 3). This shows that QPM.FO_1D.2 mediates PM resistance only at the adult plant stage and thereby confirms its role in APR.

**Figure 3.**
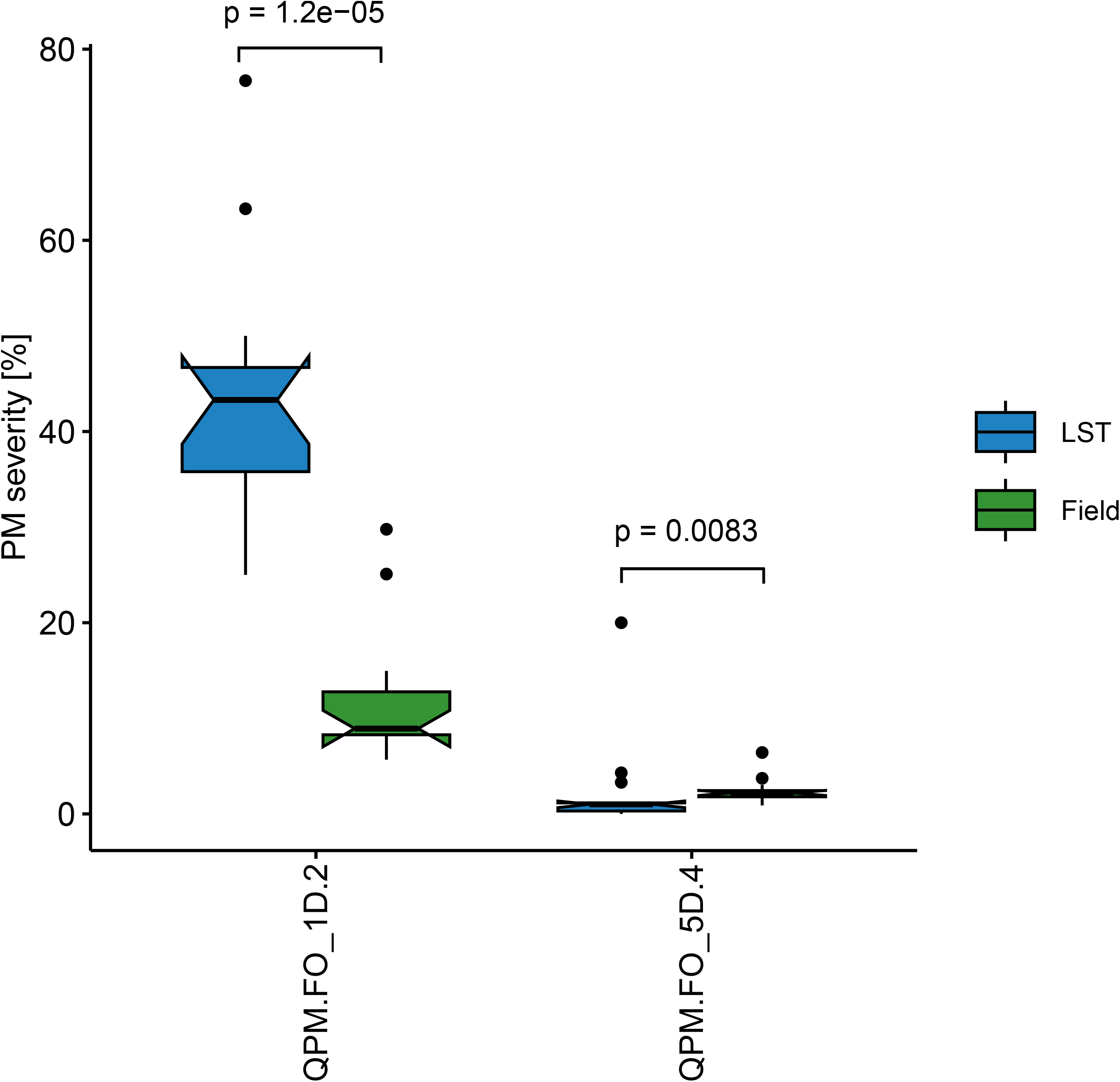
Powdery mildew severity [%] of oat lines carrying the respective favourable allele at the identified QTL on chromosome 1D (QPM.FO_1D.2) and 5D (QPM.FO_5D.4; Pm7) in leaf-segment test (LST) and in the field. QPM.FO_1D.2 includes genotypes that carry the favourable allele at avgbs2_84146.1.21 and QPM.FO_5D.4 includes genotypes that carry the favourable allele at the significant MTA for Pm7 (avgbs_102450.1.35). Kruskal-Wallis p values are given above the brackets.

### QTL mapping

The oat variety Husky is a close relative of Firth, in which the PM APR gene on chromosome 1D was first described (Hagmann et al., 2012). Hence, to further validate the location of the APR gene, a QTL mapping experiment with 176 lines from a HuskyxAVE1284 cross was performed. AVE1284 is an oat land race that is moderately susceptible to PM infection. The population comprised two subpopulations that each consisted of 101 (subpopulation 1) and 75 (subpopulation 2) individuals, respectively. Assessment of PM severity revealed that PM resistance only segregated in the first, but not in the second subpopulation (Supplementary Figure S3). Hence, subpopulation 2 was excluded from all subsequent analyses. Mean PM severity in the population across all environments was 14.0 %, with the majority of observations being distributed between the parental means (Husky: 5.1 % and AVE1284: 20.4 %), however values outside that range were observed as well (Supplementary Table S1, Figure 3A). Broad-sense heritability of logit transformed PM severity was estimated to be 87.0 %, reflecting a good reproducibility of observations across environments (Supplementary Table S1).

GBS calls filtered at 50% completeness provided genotypes for 1534 tag-level haplotypes. The relatively small number of filtered loci compared to other GBS experiments reflects the complexity reduction provided by the Rapture method. These loci were distributed across all 21 chromosomes, as was also the intention of the Rapture method. After phasing, removal of outliers, and imputation of missing genotypes, 1488 tag-level loci remained. Each subpopulation segregated on approximately 50% of chromosomes, with remaining chromosomes being fixed for the Husky allele. The first subpopulation segregated across all of chromosome 1D, while the second was fixed for Husky alleles on chromosome 1D, supporting the earlier phenotypic observations. Using the filtered and imputed markers, genetic maps could be constructed for chromosomes 1A, 1D, 2C, 2D, 3C, 4D, 5A, 5C, 5D, 6C, 6D, 7C and 7D using Joinmap (Supplementary Figure S4). The genetic map comprised 527 markers and spanned across 471 cM, giving an average marker spacing of 0.89 cM. Due to limited recombination in the rather small population, maps could not be constructed for all chromosomes and two chromosomes (3C and 4D) are represented by two separate linkage groups. Correlation analysis between the genetic and physical marker order showed good consistency between the genetic map and the Sang reference genome (Spearman rank correlation 0.99, (Kamal et al., 2022)).

QTL detection identified a single significant QTL region at the distal end of chromosome 1D, designated QPM.HuskyxAVE1284_1D (α < 0.05, Figure 4B). The LOD score of the most significant marker was 22.44 and the percentage of phenotypic variance explained by the QTL was 64 %. In individuals carrying the Husky allele (AA) at the identified marker, PM severity was reduced compared to individuals carrying the AVE1284 allele (BB). Individuals that were heterozygous at the identified marker (AB) showed intermediate PM severity, suggesting an additive positive effect on PM resistance of the Husky allele (Figure 4C). The physical region of the QTL spanned 27.32 Mb and contained 573 high-confidence genes based on the Sang oat genome assembly (Position 442676872 to 469999753; Kamal et al., 2022). Within these, several genes were predicted to encode disease resistance proteins such as NLRs (nucleotide-binding leucine-rich repeat receptors; 15 genes) and CRKs (cysteine-rich receptor-like kinases; 5 genes). Furthermore, two genes in this region were homologs of the barley PM susceptibility gene MLO (Mildew-Locus-O, Jørgensen, 1992). In addition, eight genes were predicted to belong to the protein kinase superfamily.

**Figure 4.**
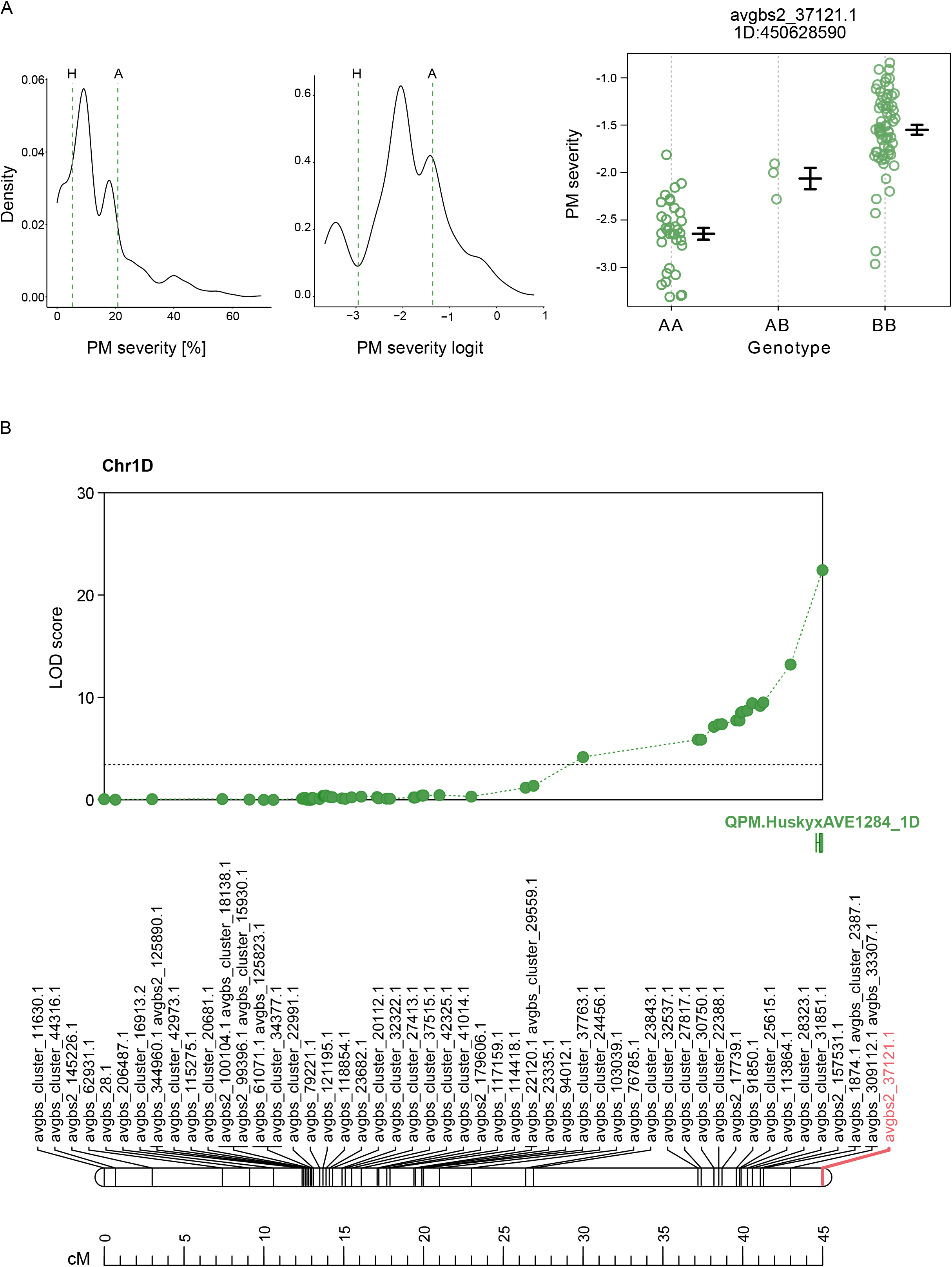
QTL mapping for powdery mildew resistance. **A**. Distribution of powdery mildew severity [%] (left) and logit-transformed values (right). Means of parental individuals are indicated by dashed vertical lines. H: Husky; A: AVE1284. **B**. Genetic linkage map of chromosome 1D with significant QTL identified in the HuskyxAVE1284 population. The identified most significant marker is highlighted in red. Genetic distances are indicated in centimorgans (cM) at the bottom of the chromosome. A LOD threshold of 3.43 calculated through 1000 permutations with α=0.05 is shown by a dotted vertical line. **C**. Phenotype distribution for the genotypes of the identified significant marker with logit-transformed values for powdery mildew severity. AA refers to the genotype from Husky, BB to the genotype from AVE1284, and AB to the heterozygous genotype.

### Comparison of GWAS and QTL mapping results

Comparison of the position of the significant markers from GWAS with the QTL identified in the HuskyxAVE1284 population did not show an overlap, but the two more terminal significant regions QPM.FO_1D.2 and QPM.FO_1D.3 were fairly close to QPM.HuskyxAVE1284_1D (2.2 and 24.1 Mb distance, Supplementary Figure S5). Furthermore, by analysis of linkage disequilibrium (LD) we identified two distinct large linkage blocks in the genetic map of chromosome 1D, indicating limited recombination within these blocks (Supplementary Figure S5). The second block spans from position 383744256 to the end of the chromosome and contains QPM.HuskyxAVE1284_1D as well as the two terminal significance regions QPM.FO_1D.2 and QPM.FO_1D.3 from GWAS.

Taken together, both approaches independently positioned the PM APR gene on the distal end of chromosome 1D, being in agreement with and further refining the position previously described by Hagmann et al. (2012). The locus was confirmed to confer APR and the underlying gene is designated *Pm13* to be consistent with the oat PM resistance gene nomenclature and latest described PM gene (Hsam et al., 2014; Ociepa et al., 2022). For more precise positioning of the gene, an additional QTL mapping approach using a larger population should be considered.

## Discussion

The aim of this study was to narrow down the location of the powdery mildew APR gene present in the related oat varieties Firth, Flämingstip and Husky. For this, two separate experiments were conducted. One experiment involved Husky and employed linkage mapping in a bi-parental population, while the other approach used GWAS in a population including Firth and related breeding strains.

Linkage mapping and GWAS both positioned QTLs for powdery mildew severity on the distal end of chromosome 1D, providing strong evidence for the location of the APR and being in line with the results of Hagmann et al. (2012). However, although they were quite close, there was no perfect overlap between QTLs identified in GWAS and QTL mapping. This could be due to the small population size used for the QTL analysis, as genetic maps based on dominant markers in small populations are prone to mis-ordering (Ferreira et al., 2006). Analysis of linkage disequilibrium identified two linkage blocks with reduced recombination on chromosome 1D (Supplementary Figure S5). Two of the QTLs identified by GWAS lie within the second linkage block, which also contains the QTL identified by QTL mapping. Strong marker linkage may mean that the actual causal locus is not within the identified QTL, but at a locus within this linkage block. This could further explain the discrepancy between QTL mapping and GWAS results. Another explanation could be that Husky from the QTL study and the related varieties Firth, Flämingstip etc. from the GWAS study have different PM resistance genes. The present results do not exclude this possibility. However, given the close relationship of the mentioned oat varieties and proximity of the identified QTLs, it is more probable that they are linked to the same gene. Hence, we consider it sufficiently likely that the APR gene in Husky, Firth and other related varieties is the same gene, that we tentatively designate Pm13. More precise mapping and marker development seems necessary to validate the results. Furthermore, different statistical models in GWAS often lead to different MTAs being identified, as was the case in our study. This further highlights the importance of validating associations identified by GWAS.

In addition to Pm13, other PM resistance genes were also present in the FO-panel, as suggested by the identification of additional significant MTAs on chromosomes 1A, 5D, 6C, 7A and 7D. The MTAs on chromosome 1A, 5D and 7A can be attributed to the resistance genes Pm3 (Mohler, 2022), Pm7 (Brodführer et al., 2023) and Pm11 (Ociepa et al., 2020), respectively. To the best of our knowledge, the MTAs on 6C and 7D could be linked to new, unmapped resistance loci.

As opposed to race-specific resistance, it is generally accepted that APR is more durable and thought to be under polygenic control. However, a single main QTL associated with APR was identified in this study despite the continuous variation in the degree of infection found. GWAS revealed the possibility of additional minor QTL and further analysis is required to determine their role. Conversely APR can be viewed as an incomplete and delayed protection of a plant against a pathogen and could be governed by a single gene as suggested by this study. APR resistance genes do not provide absolute resistance as conferred by major genes such as Pm7 but partial resistance. APR against mildew has been reported in both wheat and barley and have been found to be both single or polygenic (Asad et al., 2014; Burdon et al., 2014; Johnson et al., 2003) and may even provide resistance to more than one fungal pathogen, for example Lr34/Yr18/Pm38 and Lr67/Yr46/Sr55/Pm46 in wheat (Herrera-Foessel et al., 2014; Krattinger et al., 2009).

In a number of cases, kinases have been shown to play an important role in APR. For example, two maize wall-associated kinases, ZmWAK-RLK1 and ZmWAK, confer APR to Northern leaf blight and head smut, respectively (Hurni et al., 2015; Zuo et al., 2015). In accordance with this, the most promising candidate gene from GWAS is predicted to encode a protein kinase superfamily protein. Furthermore, eight genes within the QTL from QTL mapping are also predicted to be members of this gene family. Several other genes in the QTL region from the QTL mapping are predicted to encode NLRs or CRKs. Even though NLRs and CRKs are typical qualitative R genes, they can also be involved in APR (Barbacci et al., 2020; Poland et al., 2009). Furthermore, two MLO-like genes are located within the QTL from QTL mapping. MLO is a powdery mildew susceptibility gene that was initially identified in barley but that is broadly conserved across plant species (Appiano et al., 2015; Jørgensen, 1992). In oats, several MLO-like genes have been identified based on sequence homologies (Reilly et al., 2021), but their functionality has not been shown yet. Taken together, even though several intriguing putative functions of Pm13 have been identified, they remain speculative and additional efforts are necessary to fine map and clone Pm13.

Strikingly, we observed significantly lower PM severity in the autumn-sown trials than in the spring-sown trials at all locations. PM infection is strongly influenced by weather conditions such as temperature, humidity and rainfall. The most favourable temperature for infection is approximately 15–20°C, and spore germination is facilitated by high relative humidity, but the presence of free water hinders spore germination (Agrios, 2005; Cowger et al., 2012). Hence, the lower temperatures and higher rainfalls in winter could explain our observation. In wheat and barley, the effect on sowing date on PM infection has been studied as well. There, later sowings also resulted in higher disease incidences compared to earlier sowings (Last, 1957; Naseri & Sheikholeslami, 2021). Similarly, Reilly et al. (2024) evaluated PM resistance in autumn and spring seeded experiments of the oat cultivars Husky and Keely as well as Pm7 carriers Delfin, Elison and Yukon. Interestingly, there, autumn sown susceptible oats were higher infected than the spring sown, which is in contrast with our observations. This is surprising, because the lower BGA infection of the autumn sown variant in our study was consistent also in Aberystwyth, with similar climatic conditions as in Ireland of the, Reilly et al. (2024) study. However, overall infection levels in, Reilly et al. (2024) were much lower than in our study, with maximum values of around 8 % infected leaf area. Differences in disease pressure might be the reason for our contrasting observations. In addition, Reilly et al. (2024) found a significantly higher BGA incidence in Husky compared to Keely in some environments. This may be related to an adaptation of BGA to Pm13 in Husky, due to many years of growing of Husky on a large area in Ireland. It is very interesting that an adaptation of BGA to a single resistance QTL in one cultivar might be possible even though a related cultivar with the same QTL is not affected in the same way. This resembles the overcome of Pm7 in the cultivars Canyon, Delfin and Yukon by polish BGA isolates (Reilly et al., 2024), but without any natural selections, since these Pm7 carriers were not widely grown in those areas in Poland the BGA isolates came from. Natural BGA virulence spectrum seems to be high and even the Pm13 gene seems not to be immune to degradation. Also in wheat, differences in PM severity between cultivars carrying the same Pm resistance gene have been reported, which was attributed to the presence of additional minor resistance loci in some of these cultivars (Miedaner et al., 2007). This could also be the case in oats and could explain these observations.

However, when looking at the differences between Husky and Keely in Reilly et al. (2024), the overall low levels of infection must be taken into account. Although statistically significant, their functional importance can be questioned due to the low magnitude of differences.

In conclusion, the combination of linkage mapping and GWAS provided valuable insights into the location of the powdery mildew APR gene Pm13 within the closely related oat varieties Firth, Flämingstip, and Husky, pointing to the distal end of chromosome 1D as the likely location. The presence of kinases, NLRs, CRKs, and MLO-like genes in the QTL regions suggests potential candidates for further exploration. Additionally, the observed seasonality effects on PM severity underscore the influence of environmental conditions on disease dynamics.

## Funding

This study is part of the CROPDIVA project “Climate Resilient Orphan croPs for increased DIVersity in Agriculture” funded by the EU’s Horizon 2020 Research and Innovation Programme (Grant 364 Agreement No. 101000847).

## Footnotes

1 https://www.indexmundi.com/agriculture/?country=eu&commodity=oats&graph=production

## Notes

### Competing Interest Statement

The authors have declared no competing interest.

