## Supplementary for "Genetic mapping of the powdery mildew resistance gene Pm13 on oat chromosome 1D"

**Supplementary Figure S1: Phylogenetic tree of oat lines in the KLAR panel.** The closely related oat lines Husky, Keely, Firth, Flämingstip and Winston are highlighted in yellow. Markers were filtered for a minor allele frequency of 0.05 and 95 % completeness. Marker data extracted from Brodführer et al. (2023).

**Supplementary Figure S2.** Phenotype distribution of the genotypes for significant SNP markers detected in GWAS. Marker names with their respective chromosomal position are given at the top. Genotypes are given in IUPAC code.

**Supplementary Figure S3. Powdery mildew severity of HuskyxAVE1284 subpopulations in three environments.** GL.: Groß Lüsewitz. QLB: Quedlinburg.

**Supplementary Figure S4: Genetic map for the HuskyxAVE1284 population based on Rapture-GBS markers.** Black lines indicate marker positions and colours along the chromosomes indicate marker density. Figure rendered using LinkageMapView 2.1.2 (Ouellette *et al.*, 2018).

**Supplementary Figure S5: Linkage of markers of the genetic map of chromosome 1D for the HuskyxAVE1284 population and relative positions of loci identified by GWAS and QTL mapping.** Color of the square at the intersection of two loci represents their linkage (R2; red and pink depict loci with high linkage, blue and white depict loci with low linkage). Grey bars on top represent relative positions of the QTLs identified by GWAS (QPM.FO_1D.1, QPM.FO_1D.2, QPM.FO_1D.3) and QTL mapping (QPM.HuskyxAVE1284_1D). Linkage was analysed in TASSEL. Size of loci not to scale.

**Supplementary Table S1. Means and variance components of PM severity data in the FO-panel and HuskyxAVE1284 (HxA) population.** Mean, minimum and maximum values are given for the untransformed data, variance components are calculated based on the logit-transformed values.

| | **Mean** | **Min** | **Max** | **$LSD_{5\%}$** | **$Var.cp_G$** | **$Var.cp_{GxE}$** | **$Var.cp_E$** | **$h^2$** |
|---|---|---|---|---|---|---|---|---|
| **FO (standards only)** | 18.76 | 0 | 90 | 2.29 | 3.933 | 0.104 | 2.103 | 88.54 |
| **FO entries** | 22.3 | 0 | 100 | 0.82 | 0.43 | 0.10 | 0.86 | 83.13 |
| **HxA** | 13.97 | 0 | 70 | 0.65 | 0.3616 | 0.0912 | 0.2623 | 87.05 |

**Supplementary Table S2. Flanking sequence with SNPs of significant markers identified by GWAS and QTL mapping on chromosome 1D.** SNPs are highlighted in bold. Chr.: Chromosome. Pos.: Position in the Sang genome assembly. QTL: Name of the QTL the marker was assigned to.

| **Marker** | **Position** | **QTL** | **Flanking sequence with SNP** |
|---|---|---|---|
| **avgbs_cluster_38613.1.56** | 343715080 | QPM.FO_1D.1 | TGCAGTGGCAGGCATCCTYGAAACAGGCTTTCGCCCCGCTATATTAATATAGCAG**[C/T]**GGCAGGCA |
| **avgbs2_84146.1.21** | 418134649 | QPM.FO_1D.2 | TGCAGCGAGAAAGACGTAAG**[C/T]**ATCATCCGTGGCTGCTGGCGTCGCCTTGCTGTGTTATGATGGC |
| **avgbs_47533.1.46** | 418137530 | QPM.FO_1D.2 | TGCAGCCGATCCTCCAGTTTCGATTGACGGACAGCTTCCTAATTC**[C/G]**TTTTTCGACTTCTCTTCT |
| **avgbs_69900.1.31** | 440373230 | QPM.FO_1D.3 | TGCAGCTCCGCTCTCGGCGCCGACCAGTTC**[A/G]**ACGACGCCGATCCAGTGGAGCGGGTGGCCAAGG |
| **avgbs2_37121.1** | 450628590 | QPM.HuskyxAVE1284_1D | TGCAGATGGTTTTGTGAAGCTACATCAAAAC**[C/T]TCAGAAGT[C/T]AC[C/T]**GCATACTTTGGCTGTCACAA |

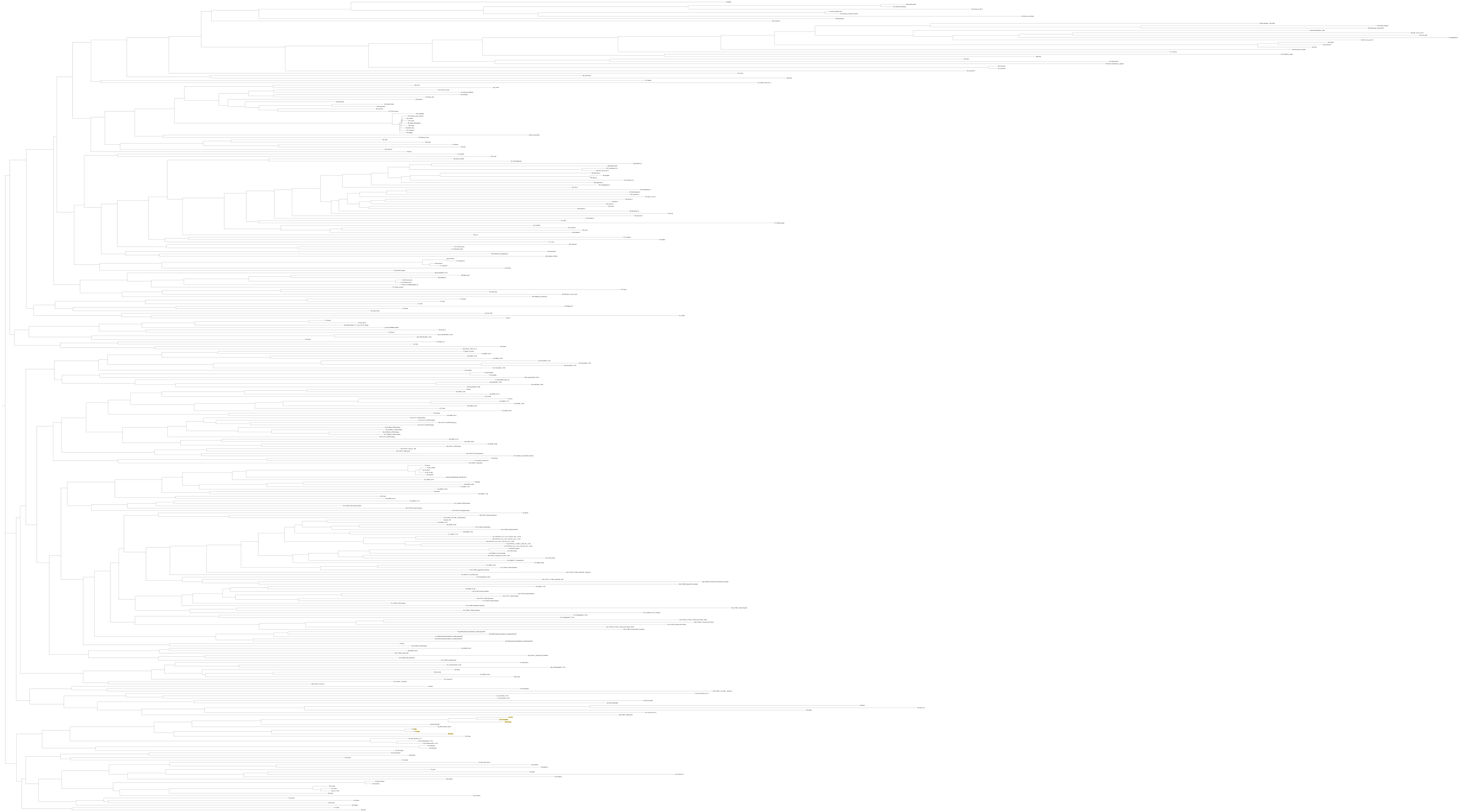

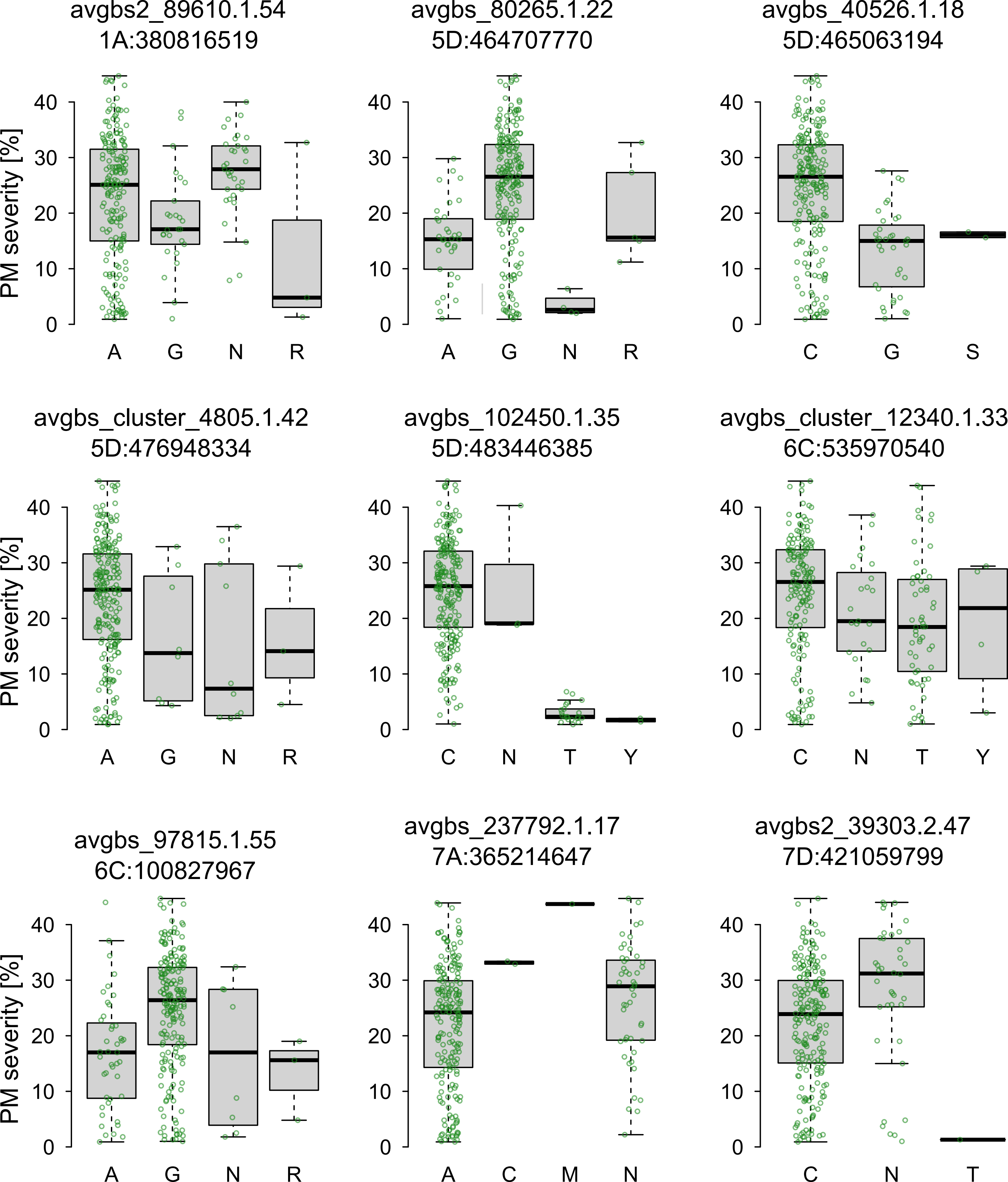
avgbs2_89610.1.54
1A:380816519
avgbs_80265.1.22
5D:464707770
avgbs_40526.1.18
5D:465063194
avgbs_cluster_4805.1.42
5D:476948334
avgbs_102450.1.35
5D:483446385
avgbs_cluster_12340.1.33
6C:535970540
avgbs_97815.1.55
6C:100827967
avgbs_237792.1.17
7A:365214647
avgbs2_39303.2.47
7D:421059799
PM severity [%]
0
10
20
30
40
A
G
N
R
C
S
T
Y
M

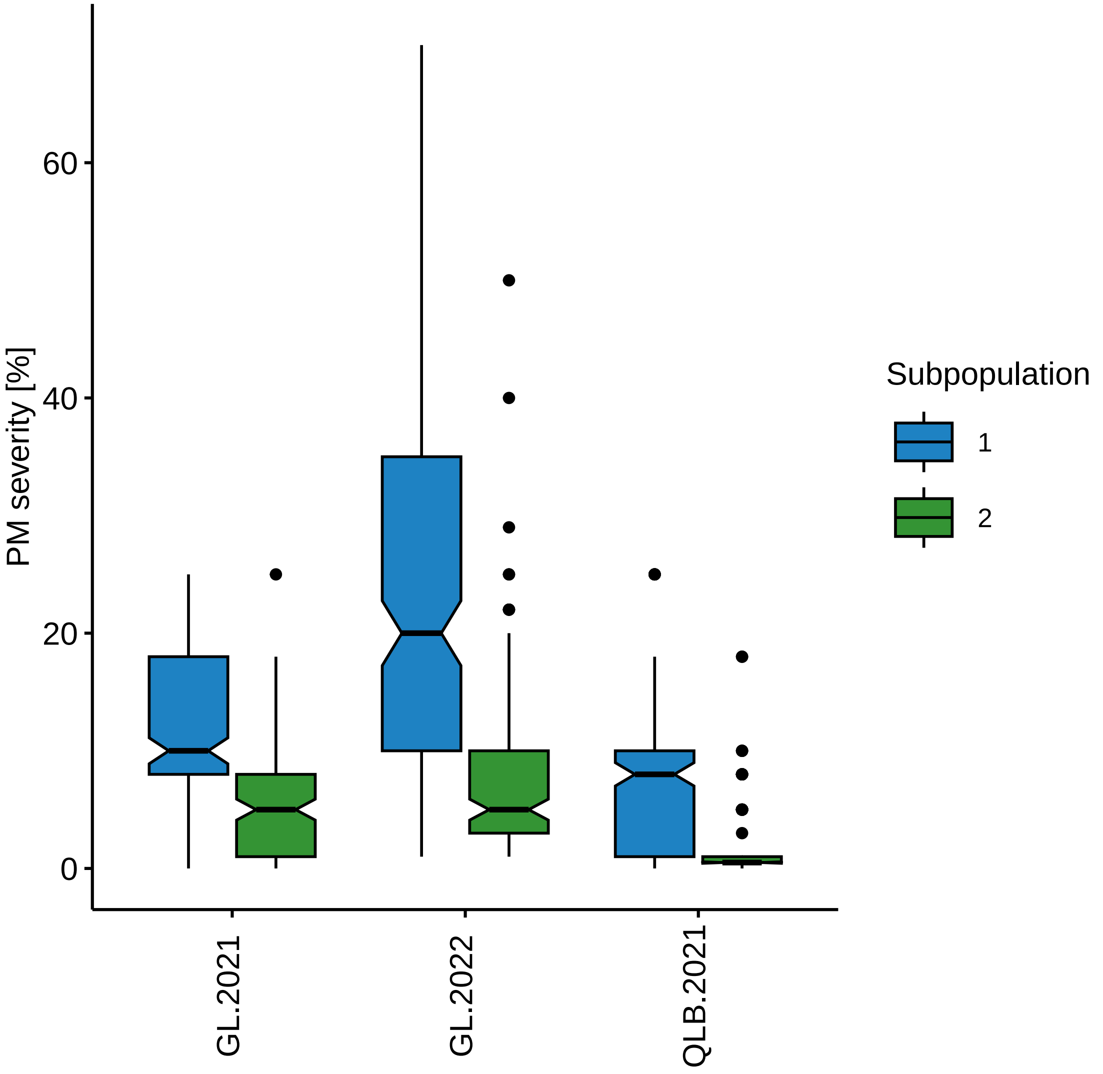

PM severity [%]
60
40
20
0
GL.2021
GL.2022
QLB.2021
Subpopulation
1
2

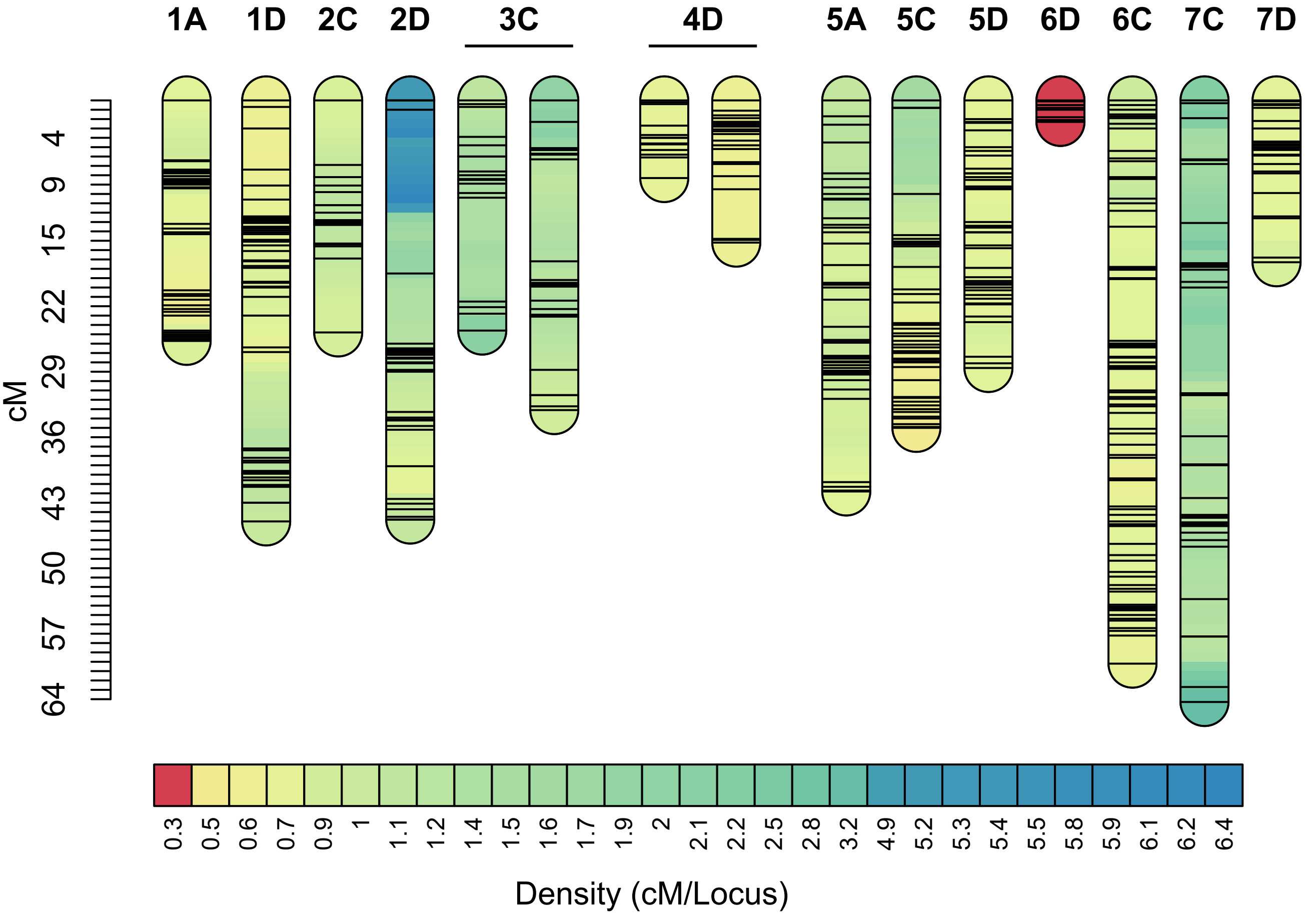
1A 1D 2C 2D 3C 4D 5A 5C 5D 6D 6C 7C 7D
cM
4 9 15 22 29 36 43 50 57 64
0.3 0.5 0.6 0.7 0.9 1 1.1 1.2 1.4 1.5 1.6 1.7 1.9 2 2.1 2.2 2.5 2.8 3.2 4.9 5.2 5.3 5.4 5.5 5.8 5.9 6.1 6.2 6.4
Density (cM/Locus)

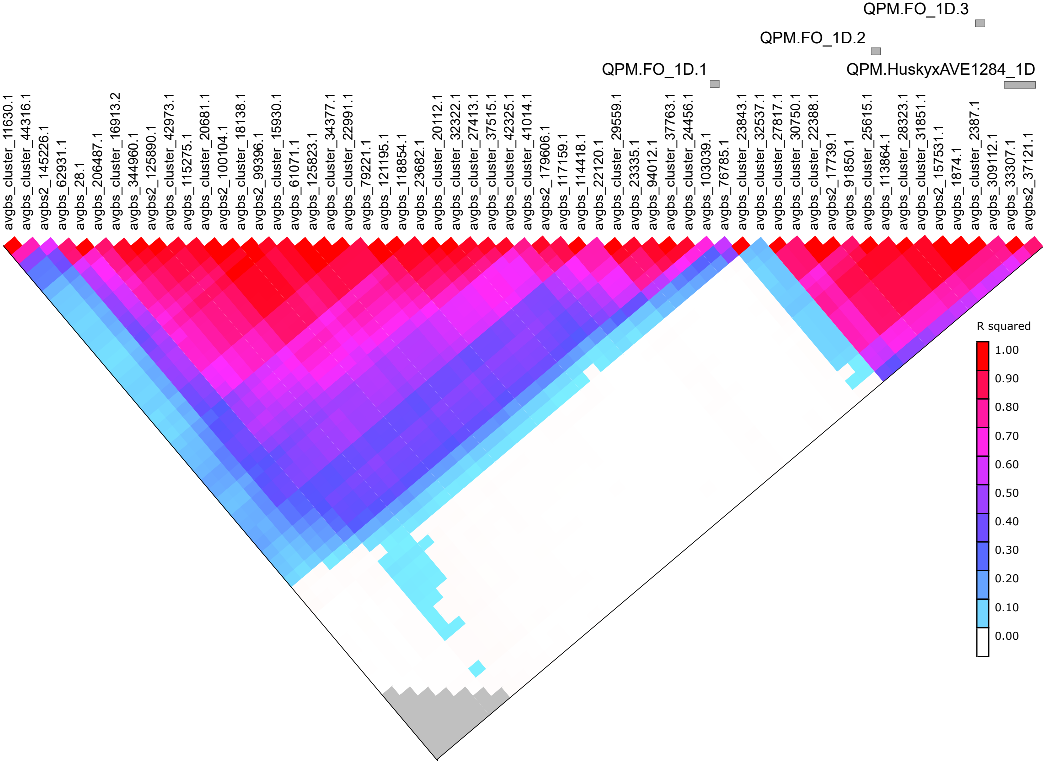
QPM.FO_1D.3
QPM.FO_1D.2
QPM.FO_1D.1
QPM.HuskyxAVE1284_1D
avgbs_cluster_11630.1
avgbs_cluster_44316.1
avgbs2_145226.1
avgbs_62931.1
avgbs_28.1
avgbs_206487.1
avgbs_cluster_16913.2
avgbs_344960.1
avgbs2_125890.1
avgbs_cluster_42973.1
avgbs_115275.1
avgbs_cluster_20681.1
avgbs2_100104.1
avgbs_cluster_18138.1
avgbs2_99396.1
avgbs_cluster_15930.1
avgbs_61071.1
avgbs_125823.1
avgbs_cluster_34377.1
avgbs_cluster_22991.1
avgbs_79221.1
avgbs_121195.1
avgbs_118854.1
avgbs_23682.1
avgbs_cluster_20112.1
avgbs_cluster_32322.1
avgbs_cluster_27413.1
avgbs_cluster_37515.1
avgbs_cluster_42325.1
avgbs_cluster_41014.1
avgbs2_179606.1
avgbs_117159.1
avgbs_114418.1
avgbs_22120.1
avgbs_cluster_29559.1
avgbs_23335.1
avgbs_94012.1
avgbs_cluster_37763.1
avgbs_cluster_24456.1
avgbs_103039.1
avgbs_76785.1
avgbs_cluster_23843.1
avgbs_cluster_32537.1
avgbs_cluster_27817.1
avgbs_cluster_30750.1
avgbs_cluster_22388.1
avgbs2_17739.1
avgbs_91850.1
avgbs_cluster_25615.1
avgbs_113864.1
avgbs_cluster_28323.1
avgbs_cluster_31851.1
avgbs2_157531.1
avgbs_1874.1
avgbs_cluster_2387.1
avgbs_309112.1
avgbs_33307.1
avgbs2_37121.1
R squared
1.00
0.90
0.80
0.70
0.60
0.50
0.40
0.30
0.20
0.10
0.00
